# ReferenceSeeker: rapid determination of appropriate reference genomes

**DOI:** 10.1101/863621

**Authors:** O. Schwengers, T. Hain, T. Chakraborty, A. Goesmann

**Author notes:** Contributed equally to this work (T. Chakraborty, A. Goesmann).

## Abstract

**Summary:** The large and growing number of microbial genomes available in public databases makes the optimal selection of reference genomes necessary for many in-silico analyses, e.g. single nucleotide polymorphism detection, scaffolding and comparative genomics, increasingly difficult. Here, we present ReferenceSeeker, a novel command line tool combining a fast kmer profile-based database lookup of candidate reference genomes with subsequent calculation of highly specific average nucleotide identity (ANI) values for the rapid determination of appropriate reference genomes. Pre-built databases for bacteria, archaea, fungi, protozoa and viruses based on the RefSeq database are provided for download.

**Availability and Implementation:** ReferenceSeeker is open source software implemented in Python. Source code and binaries are freely available for download at https://github.com/oschwengers/referenceseeker under the GNU GPL3 license.

## Introduction

The enormous success and ubiquitous application of next and third generation sequencing has led to a large number of available high-quality draft and complete microbial genomes in the public databases. Today, the NCBI RefSeq database release 90 alone contains 11,060 complete bacterial genomes (Haft et al. 2018). Concurrently, selection of appropriate reference genomes (RGs) is increasingly important as it has enormous implications for routine in-silico analyses, as for example in detection of single nucleotide polymorphisms, scaffolding of draft assemblies, comparative genomics and metagenomic tasks. Therefore, a rigorously selected RG is a prerequisite for the accurate and successful application of the aforementioned bioinformatic analyses. In order to address this issue several new databases, methods and tools have been published in recent years e.g. RefSeq, DNA-DNA hybridization (Meier-Kolthoff et al. 2013), average nucleotide identity (ANI) values (Goris et al. 2007) and Mash (Ondov et al. 2016). Nevertheless, the sheer amount of currently available databases and potential RGs contained therein, together with the plethora of tools available, often require manual selection of the most suitable RGs. To the best of the authors’ knowledge, there is currently no such tool providing both an integrated, highly specific workflow and scalable and rapid implementation. ReferenceSeeker was designed to overcome this bottleneck. As a novel command line tool, it combines a fast kmer profile-based lookup of candidate reference genomes (CRGs) from high quality databases with rapid computation of highly specific ANI and conserved DNA values.

## Implementation

ReferenceSeeker is a linux command line tool implemented in Python 3. All necessary external binaries are bundled with the software. The tool Itself requires no external dependencies other than Biopython for file input and output.

### Databases

ReferenceSeeker takes advantage of taxon-specific custom databases in order to reduce data size and overall runtime. Pre-built databases for the taxonomic groups bacteria, archaea, fungi, protozoa and viruses are provided. Each database integrates genomic as well as taxonomic information comprising genome sequences of all RefSeq genomes with an assembly level ‘complete’ or whose RefSeq category is either denoted as ‘reference genome’ or ‘representative genome’, as well as kmer profiles, related species names, NCBI Taxonomy identifiers and RefSeq assembly identifiers. For convenient and fully automatic updates, we provide locally executable scripts implemented in bash and Nextflow (Di Tommaso et al. 2017).

### Database Lookup of CRGs

To reduce the number of necessary ANI calculations a kmer profile-based lookup of CRGs against custom databases is carried out. This step is implemented via Mash parameterized with a Mash distance of 0.1, which was shown to correlate well with an ANI of roughly 90% (Ondov et al. 2016) and thereby establishing a lower limit for reasonably related genomes. The resulting set of CRGs is subsequently reduced to a configurable number of CRGs (default=100) with the lowest Mash distances.

### Determination of RG

As a highly specific measure for microbial genome relationships ReferenceSeeker uses ANI and conserved DNA (conDNA) values. The reasoning for the subsequent calculation of ANI and conDNA values is that although Mash distance values correlate well with ANI values, the same cannot be observed for conDNA values (Fig. 1).

**Figure 1.**
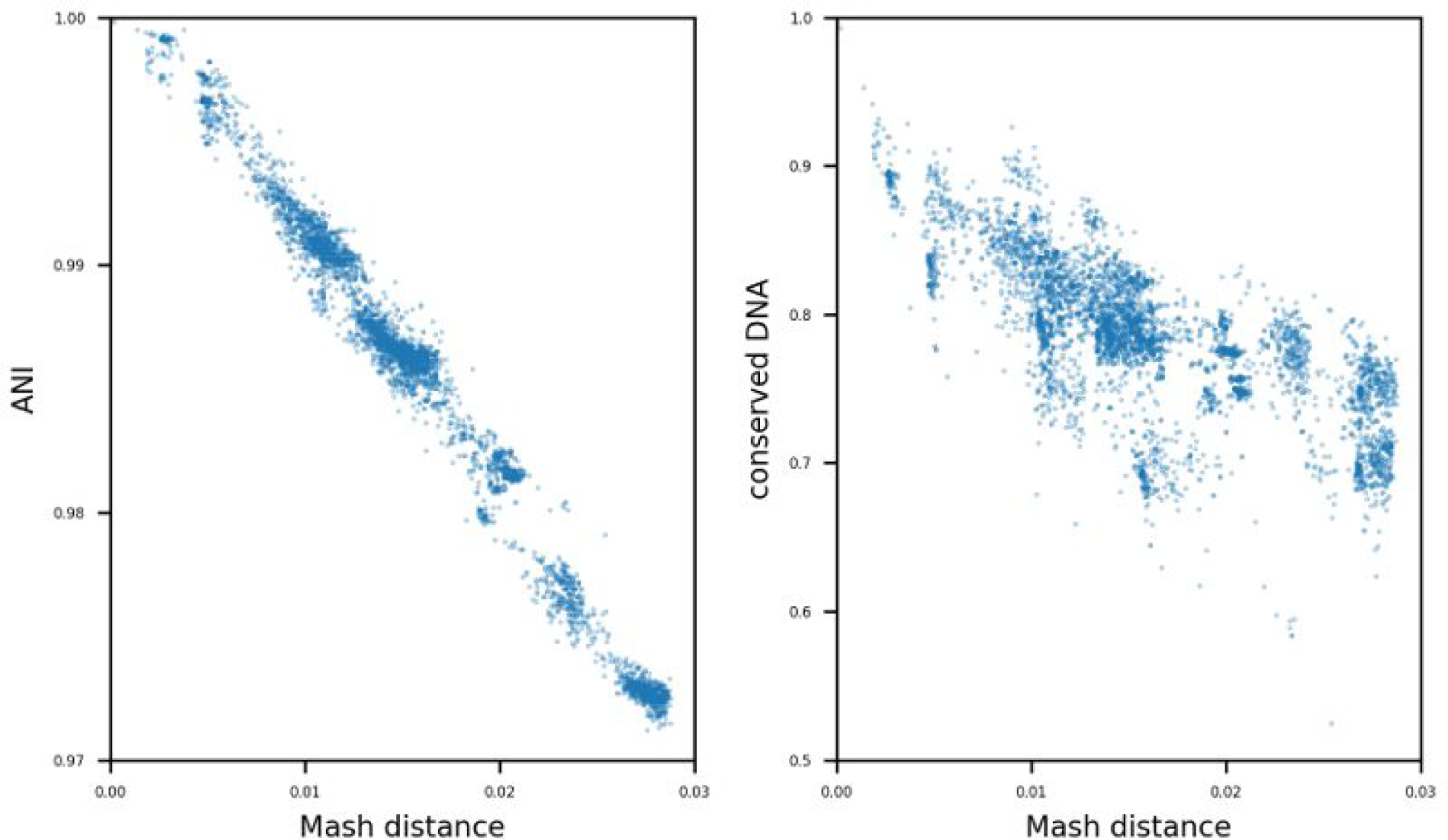
Scatter plots showing the correlation between Mash distance, ANI and conDNA values. ANI and conserved DNA values are plotted against Mash distance values for 500 candidate reference genomes with the lowest Mash distance within the bacterial database for 10 randomly selected *Escherichia coli* genomes from the RefSeq database, each.

Calculation of these metrics is implemented vía Nucmer contained in the MUMmer package (Marçais et al. 2018) as it was recently shown that Nucmer based implementations (ANIn) compare favourably against BLAST+ based implementations (ANIb) in terms of runtime. Given that compared genomes are closely related, i.e. shared ANI is above 90%, it was also shown that ANIn correlates well with ANIb (Yoon et al. 2017). This is ensured by the prior Mash-based selection of CRGs. As an established threshold for species boundaries (Goris et al. 2007), results are subsequently filtered by configurable ANI and conDNA values with a default of 95% and 69%, respectively. Finally, CRGs are sorted according to the harmonic mean of ANI and conDNA values in order to incorporate both nucleotide identity and genome coverage of query genomes and resulting CRGs. In this manner, ReferenceSeeker ensures that the resulting RGs sufficiently reflect the genomic landscape of a query genome.

### Application

ReferenceSeeker takes as input a microbial genome assembly in fasta format and the path to a taxonomic database of choice. Results are returned as a tabular separated list comprising the following information: RefSeq assembly identifier, ANI, conDNA, NCBI taxonomy identifier, assembly status and organism name.

To illustrate the broad applicability at different scales we tested ReferenceSeeker with 12 microbial genomes from different taxonomic groups and measured overall runtimes on a common consumer laptop providing 4 cores and a server providing 64 cores (Table 1). For the tested bacterial genomes, ReferenceSeeker limited the number of resulting RGs to a default maximum of 100 genomes. Runtimes of archaeal and viral genomes are significantly shorter due to a small number of available RGs in the database and overall smaller genome sizes, respectively.

**Table 1.** Runtimes and numbers of resulting RG executed locally on a quad-core moderate consumer laptop and a 64 core server machine.

| Genome | Genome Size<br>[kb] | Laptop<br>[m:s] | Server<br>[m:s] | # RG |
| --- | --- | --- | --- | --- |
| *Escherichia coli* str. K-12 substr. MG1665<br>(GCF_000005845.2) | 4,641 | 3:24 | 0:30 | 100* |
| *Pseudomonas aeruginosa* PAO1<br>(GCF_000006765.1) | 6,264 | 5:20 | 0:44 | 100* |
| *Listeria monocytogenes* EGD-e<br>(GCF_000196035.1) | 2,944 | 2:52 | 0:24 | 100* |
| *Staphylococcus aureus* subsp aureus NCTC<br>8325 (GCF_000013425.1) | 2,821 | 2:31 | 0:21 | 100* |
| *Halobacterium salinarum* NRC-1<br>(GCF_000006805.1) | 2,571 | 0:04 | 0:03 | 2 |
| *Methanococcus maripaludis* X1<br>(GCF_000220645.1) | 1,746 | 0:22 | 0:09 | 5 |
| *Aspergillus fumigatus* Af293<br>(GCF_000002655.1) | 29,384 | 3:11 | 2:07 | 1 |
| *Candida albicans* SC5314<br>(GCF_000182965.3) | 14,282 | 0:21 | 0:19 | 1 |
| *Entamoeba histolytica* HM-1:IMSS<br>(GCF_000208925.1) | 20,835 | 6:04 | 4:41 | 1 |
| *Plasmodium falciparum* 3D7<br>(GCF_000002765.4) | 23,326 | 2:52 | 1:49 | 1 |
| *Influenza A virus* (GCF_001343785.1) | 13 | 0:03 | 0:02 | 1 |
| *Human coronavirus* NL63<br>(GCF_000853865.1) | 27 | 0:04 | 0:02 | 1 |

Asterisks denote number of RGs limited by ReferenceSeeker to a default maximum of 100.

## Availability

The source code is available on GitHub under a GPL3 license: https://github.com/oschwengers/referenceseeker. The software is packaged and publicly available via BioConda. Pre-built databases for bacteria, archaea, fungi, protozoa and viruses are hosted at Zenodo: https://doi.org/10.5281/zenodo.3562005. All installation instructions, examples and download links are provided on GitHub.

## ACKNOWLEDGEMENT

The authors thank Karina Brinkrolf for valuable discussions, testing and bug reports.

## REFERENCES

Di Tommaso, Paolo, Maria Chatzou, Evan W. Floden, Pablo Prieto Barja, Emilio Palumbo, and Cedric Notredame. 2017. “Nextflow Enables Reproducible Computational Workflows.” Nature Biotechnology 35 (4): 316–19. https://doi.org/10.1038/nbt.3820.

Goris, Johan, Konstantinos T. Konstantinidis, Joel A. Klappenbach, Tom Coenye, Peter Vandamme, and James M. Tiedje. 2007. “DNA-DNA Hybridization Values and Their Relationship to Whole-Genome Sequence Similarities.” International Journal of Systematic and Evolutionary Microbiology 57 (1): 81–91. https://doi.org/10.1099/ijs.0.64483-0.

Haft, Daniel H., Michael DiCuccio, Azat Badretdin, Vyacheslav Brover, Vyacheslav Chetvernin, Kathleen O’Neill, Wenjun Li, et al. 2018. “RefSeq: An Update on Prokaryotic Genome Annotation and Curation.” Nucleic Acids Research 46 (D1): D851–60. https://doi.org/10.1093/nar/gkx1068.

Marçais, Guillaume, Arthur L. Delcher, Adam M. Phillippy, Rachel Coston, Steven L. Salzberg, and Aleksey Zimin. 2018. “MUMmer4: A Fast and Versatile Genome Alignment System.” PLoS Computational Biology 14 (1): e1005944. https://doi.org/10.1371/journal.pcbi.1005944.

Meier-Kolthoff, Jan P., Alexander F. Auch, Hans-Peter Klenk, and Markus Göker. 2013. “Genome Sequence-Based Species Delimitation with Confidence Intervals and Improved Distance Functions.” BMC Bioinformatics 14: 60. https://doi.org/10.1186/1471-2105-14-60.

Ondov, Brian D., Todd J. Treangen, Páll Melsted, Adam B. Mallonee, Nicholas H. Bergman, Sergey Koren, and Adam M. Phillippy. 2016. “Mash: Fast Genome and Metagenome Distance Estimation Using MinHash.” Genome Biology 17 (1): 132. https://doi.org/10.1186/s13059-016-0997-x.

Yoon, Seok Hwan, Sung Min Ha, Jeongmin Lim, Soonjae Kwon, and Jongsik Chun. 2017. “A Large-Scale Evaluation of Algorithms to Calculate Average Nucleotide Identity.” Antonie van Leeuwenhoek, International Journal of General and Molecular Microbiology 110 (10): 1281–86. https://doi.org/10.1007/s10482-017-0844-4.

